# The lncRNAs Implicated in Redox Regulation in Ybx1 Deficient Zebrafish Larvae

**DOI:** 10.1101/679167

**Authors:** Chen Huang, Bo Zhu, Dongliang Leng, Wei Ge, Xiaohua Douglas Zhang

## Abstract

*Ybx1* has been demonstrated as a crucial gene in embryogenesis, reproduction as well as development in various vertebrates such as mouse and zebrafish. However, the underlying lncRNA-mediated mechanisms require deep investigation. Particularly, the importance of lncRNA to vertebrate development is controversial and questionable since many studies have yielded contradictory conclusions for the same lncRNAs. In the present study, in order to disclose the lncRNAs implicated in vertebrate development, a systematic transcriptome analysis is conducted based on the RNA sequencing data derived from *ybx1* homozygous mutant zebrafish on day5 (day5_*ybx1*^*-/-*^) as well as wild type zebrafish on day5 and day6 (day5_*ybx1*^*+/+*^, day6_*ybx1*^*+/+*^). A high-confidence dataset of zebrafish lncRNAs is detected using a stepwise filtering pipeline. Differential expression analysis and co-expression network analysis reveal that several lncRNAs probably act on *duox* and *noxo1a*, the genes related to redox (reduction–oxidation reaction) processes which are triggered by *ybx1* disruption. Validation by an experimental study on three selected lncRNAs indicates that knockdown of all selected lncRNAs leads to morphological deformation of larvae. In addition, our experiments effectively support the prediction of network analysis in many interaction patterns between the selected lncRNAs and the two redox genes (*duox, noxo1a*). In short, our study provides new insights into the function and mechanism of lncRNAs implicated in zebrafish embryonic development and demonstrates the importance of lncRNAs in vertebrate development.

**Author Summary:** LncRNAs has been emerged as key regulatory layers because of their multiple functions in diverse biological processes and pathways. However, there is disagreement about the roles of lncRNAs in vertebrate development Some cases demonstrated that lncRNAs was important to development. Others showed that lncRNAs just have feeble functions in development. On the other hand, Ybx1 processing multi-functions has been well demonstrated as a key protein to most vertebrate in development. Hence, aimed at disclosure of key lncRNAs implicated in vertebrate development as well as their possible roles, we performed a systematic transcriptome analysis based on the deep RNA sequencing data of *ybx1* homozygous mutant zebrafish and wild type zebrafish. Our analysis successfully revealed several lncRNAs probably target on the redox-related genes. Furthermore, our experiments validated the importance of these lncRNAs to zebrafish embryo development. This study is the first to utilize Ybx1 as breakpoint to identify key lncRNA related to vertebrate development and provides new insights into underlying mechanisms of lncRNAs in development.

## Introduction

Y-box-binding protein 1 (Ybx1, YB-1), also known as Y-box transcription factor, is a member of a large family of proteins harboring cold shock domain, and it has been proven to have multiple functions involved in a variety of biological processes and pathways, including proliferation, differentiation, regulation of apoptosis, translation, stress response, etc. This protein was found in diverse vertebrates and extensively studied in *Homo sapiens* due to its great clinical values. For instance, overexpression of Ybx1 was found in tumor cells and is associated with tumor phenotype (1). In addition, Ybx1 is demonstrated to harbor the capacity of preventing oncogenic cell transformation via the PI3K/Akt signaling pathway (2). In as much as the curial role of Ybx1 implicated in tumorigenesis, this protein has been regarded as a well-established biomarker and novel therapeutic target in cancer research (3–6).

In addition to its medicinal benefits, the multifunctional Ybx1 has also been widely utilized in the study of vertebrate development since it has been proven to be an essential gene to most vertebrate organisms for growth and development. Typically, many studies showed that *ybx1* knockout in mice could result in severe disorder in embryonic development and even death (7, 8). On the one hand, Ybx1 was regarded as a crucial protein in early embryogenesis since Ybx1-deficient mice were found to die at a very early stage of embryogenesis (8). On the other hand, it was reported that *ybx1*^−/−^embryos exhibit severe growth retardation and progressive mortality after 13.5 embryonic days, which suggests its important role in late-stage embryonic development in mice (7). Moreover, maternal Ybx1 has been demonstrated to play a safeguard to protect oocyte maturation and maternal-to-zygotic transition in zebrafish (9). The similar roles of Ybx1 were found in Pooja Kumari et al’s study (10). It was showed that Ybx1 is an essential protein to regulate the maternal control of Nodal signaling pathway which has been reported to be essential to the axis formation and germ layer specification in zebrafish (10).

In summary, Ybx1 as a key regulator in embryogenesis, reproduction as well as development has already been well demonstrated. However, the underlying mechanism of the processes and pathways mediated by Ybx1 still has a big room for deep investigation. For example, Ybx1 exerts multiple functions depending on different subcellular localizations; however, what triggers the translocation of Ybx1 from cytoplasm into nucleus has not been fully understood. Moreover, does non-coding transcriptome participate in Ybx1-mediated regulation in embryogenesis and development?

Non-coding RNA indeed has emerged in recent years as a crucial regulatory layer to participate in and govern diverse biological processes and pathways. In particular, lncRNA becomes a research hotspot in many fields since it has been demonstrated as a multifunctional regulator involved in a range of biological processes and pathways and is associated with diverse diseases (11–13). However, regarding whether lncRNA is important to vertebrate development is always debatable, particularly in embryogenesis. Mehdi Goudarzi1’s study (14) knocked down 24 specific lncRNAs in zebrafish using CRISPR/Cas9 to investigate their functions in embryo development. The result is not so satisfying as no lncRNAs are found to have overt functions in zebrafish embryogenesis, viability and fertility. Notably, a recent review by Padmanaban S. Suresha et al (15) demonstrated that a variety of non-coding RNAs mediates Ybx1 signaling, which provides a new perspective to investigate the role of lncRNA in vertebrate development. Hence, taking Ybx1 as the breakthrough to uncover the potential lncRNAs and their possible roles in embryonic development, we applied deep RNA sequencing on three different types of zebrafish larvae samples: wild type in 5 days (day5_*ybx1*^*+/+*^), wild type in 6 days (day6_*ybx1*^*+/+*^) and *ybx1*^*-/-*^ in 5 days 102 (day5_*ybx1*^*-/-*^).

Through the systematic transcriptomic analysis on our RNA sequencing data, we found several lncRNAs exhibit strong correlation to redox regulation triggered by Ybx1 knockout, i.e., response to oxidative stress, oxidation-reduction process. Furthermore, our validation experiments demonstrated the importance of three selected lncRNAs to zebrafish embryonic development and verified interaction patterns with two important Redox-related genes, i.e., *duox* and *noxo1a*. In short, our study utilized Ybx1 as a springboard to investigate the relationship of lncRNAs with zebrafish embryonic development, and successfully uncovered several relevant lncRNAs and their possible roles in it. We believe our study provides new insights into the mechanism of lncRNAs in vertebrate development.

## Results

### Identification and Characterization of lncRNAs

To detect lncRNAs in zebrafish larvae, a basic bioinformatic analysis was conducted. Briefly, the RNA sequencing yielded an average of 51,879,984 raw pair-end reads with length of 150 nt per sample (Table S1). After trimming adaptor sequences and low-quality reads, the remaining average of 49,379,818 clean reads per sample were aligned to zebrafish genome using STAR (16) and assembled into 77,252 transcripts using Stringtie (17) (Table S2). Then all the assembled transcripts were subjected to quality assessment using DETONATE (18). The results showed that the majority of transcripts has high quality. (Table S3). Additionally, BLAST search (19) against known zebrafish transcripts in Ensembl database (20) indicates that 61,926 transcripts (∼80.16%) have been well recorded in Ensembl database. Notably, the remaining 16,316 transcripts for which no hits were found in Ensembl database are probably novel transcripts representing a new subset of functional mRNAs or lncRNAs.

Based on the well-established zebrafish transcriptome, a stringent stepwise filtering pipeline was utilized to identify a high-confidence lncRNA dataset in the present study (Fig 1). Two core filtering criteria were applied to screen these lncRNAs according to the general definition of lncRNA: (1) the capability of protein-coding; (2) The length of transcripts. Briefly, 73,018 potential protein-coding transcripts obtained by BLAST search against nr, swiss-prot database as well as the known zebrafish mRNAs/proteins from Ensembl database were excluded. The remaining 1,555 unannotated transcripts shorter than 200 nucleotides in length and larger than 100 residues in the longest ORF were filtered out. A second protein-coding filtering was performed to further remove the transcripts with potential protein-coding capacity, including those containing functional domains/motifs predicted by Pfam scanning and those with protein-coding potential classified by Coding Potential Calculator (CPC) (21). This step filtered out only one transcript. A total of 2,678 lncRNAs were finally identified for further study.

**Fig 1.**
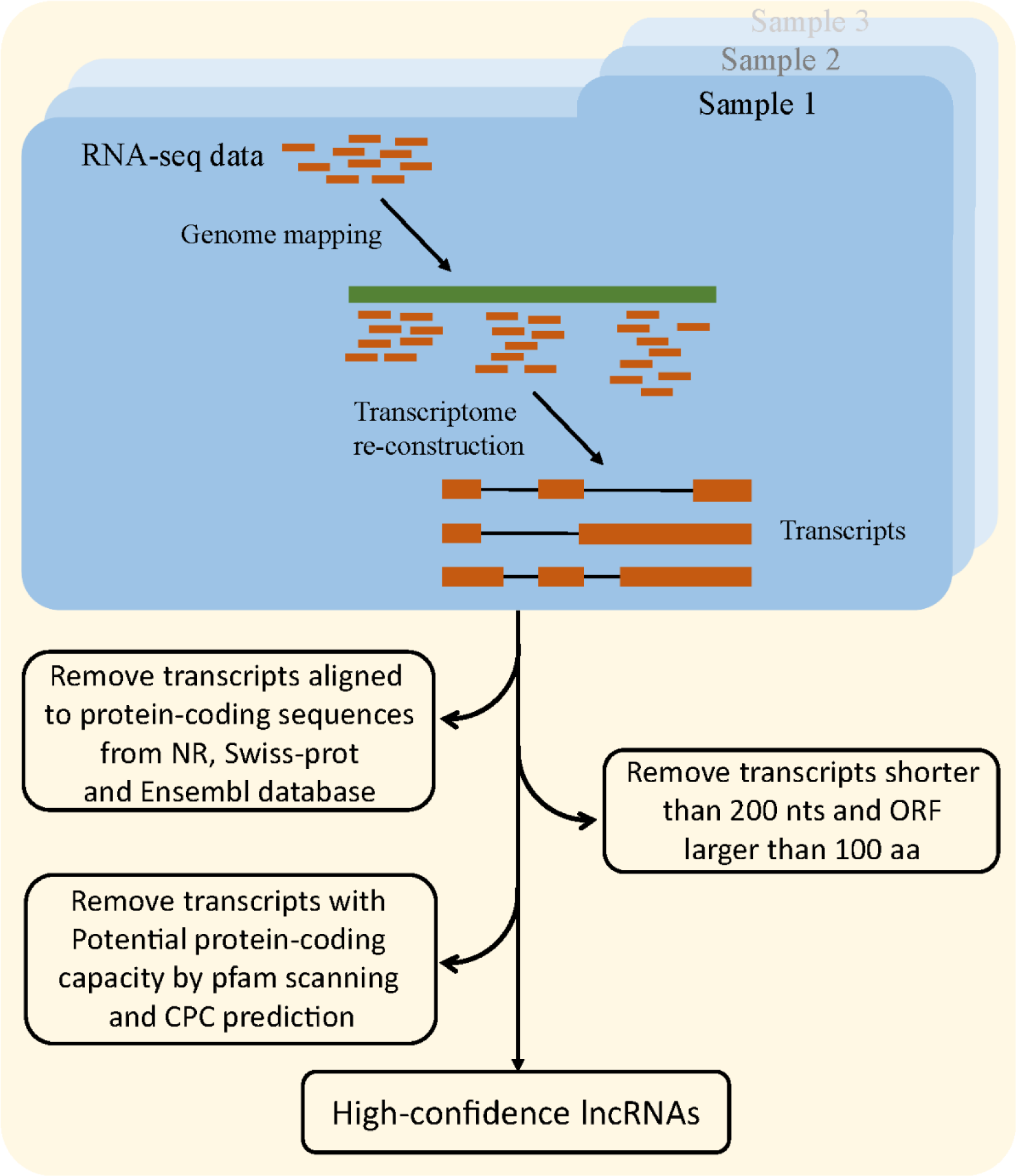
The bioinformatics pipeline utilized for identification of zebrafish lncRNAs in 778 the present study.

We compared those identified lncRNAs with known zebrafish lncRNAs derived from NONCODE database (v5.0) (22) and ZFLNC database (23), using BLASTn (e-value was set to 0.1). The results indicate that a total of 1,083 lncRNAs (∼40.44%) identified in this study could be well compared with known zebrafish lncRNAs, 460 lncRNAs of which have been included in the NONCODE and ZFLNC database. The remaining mapped lncRNAs exhibit high similarity to the known zebrafish lncRNAs in some specific regions. It is not surprising since lncRNAs have been proven to be quite distinct from mRNAs as mRNAs have more conserved mutational rate. Instead, lncRNAs have less evolutionary constraints, except the selective pressure to strictly conserve the short functional region, e.g., sequence-specific interactions, protein-binding sites, etc. The rest of lncRNAs that could not be found any hit in NONCODE and ZFLNC databases might represent a group of novel zebrafish lncRNAs. 155

### The expression pattern of the DE lncRNAs implicated in Ybx1 gene knockout

The expression profile for each sample were initially assessed using featurecounts. In order to identify the lncRNAs associated with Ybx1 gene knockout, differential expression analysis between day5_*ybx1*^*+/+*^ and day5_*ybx1*^*-/-*^ was conducted using DESeq2. The result shows that 44 lncRNAs (22 are up-regulated and 22 are down-upregulated) and 1,901 mRNAs (849 are up-regulated and 1,052 are down-upregulated) are detected as significantly differentially expressed (Fig S1, Fig S2). Despite no change of the RNA expression level of Ybx1 was observed in day6_*ybx1*^*+/+*^ compared to that of day5_*ybx1*^*+/+*^, we found that the protein expression level of Ybx1 was dramatically decreased in day6_*ybx1*^*+/+*^ vs day5_*ybx1*^*+/+*^ (data not shown). Hence, we also compared the expression level of transcripts between day5_*ybx1*^*+/+*^ and day6_*ybx1*^*+/+*^ to explore the underlying mechanisms resulting in decline of Ybx1 in day6_*ybx1*^*+/+*^ at protein level. DE analysis showed that a total of 542 RNAs were differentially expressed in day5_*ybx1*^*+/+*^ vs day6_*ybx1*^*+/+*^, including 2 DE lncRNAs and 540 DE mRNAs (Fig S3, Fig S4). Then both results of DE analysis were subjected to the DAVID server (24) to search for the possible functions of those DE RNAs. Intriguingly, both functional results indicated that the hormone synthesis-related and oxidative stress-related processes were commonly enriched with an extremely high level of statistical significance, including steroid biosynthesis process, oxidation-reduction process, etc. (Fig 2-A, B). This result disclosed a close association of Ybx1 with the processes of hormone synthesis and oxidative stress response in zebrafish.

**Fig 2.**
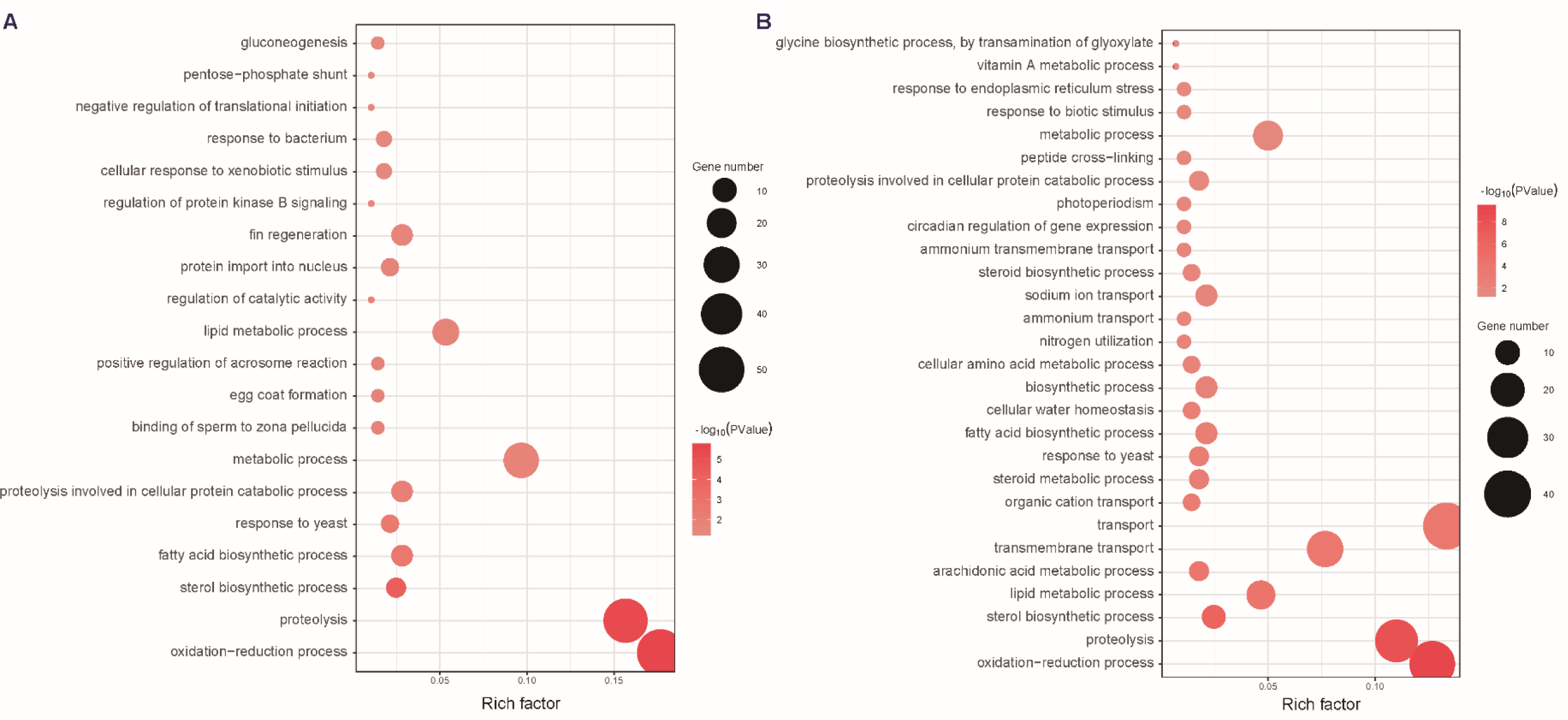
GO functional enrichment analysis of two datasets of DE transcripts achieved on 783 David server: A) day5_ybx1^+/+^ vs day5_ybx1^−/−^; B) day5_ybx1^+/+^ vs day6_ybx1^+/+^. The 784 results are only summarized in the category of biological process. The y-axis indicates 785 functional groups. The x-axis indicates –log_10_ (p value).

Also, the difference of RNA changes and the corresponding roles caused by two distinct mechanisms resulting in down-regulation of Ybx1 is another focus of attention for us to investigate. Therefore, we conducted a systematic comparison of day5_*ybx1*^*+/+*^ vs day5_*ybx1*^*-/-*^ and day5_*ybx1*^*+/+*^ vs day6_*ybx1*^*+/+*^. Concretely, 143 transcripts are found to be consistently up/down-regulated in both comparisons. Remarkably, most of them are up-regulated, and only 5 transcripts were observed as down-regulated. GO functional enrichment analysis for those 143 DE transcripts showed that many biological process (BP) categories involved in hormone-related and oxidative stress-related processes were significantly enriched, such as steroid biosynthesis, oxidation-reduction process, etc. (Fig 3). Furthermore, we found that 731 transcripts (including 22 lncRNAs) were specifically up-regulated in day5_*ybx1*^*+/+*^ vs day5_*ybx1*^*-/-*^ but exhibit no changes in day5_*ybx1*^*+/+*^ vs day6_*ybx1*^*+/+*^. Subsequent functional analysis on DAVID indicated that the functions of these specific up-regulated RNAs involved reproduction processes, i.e., egg coat formation, binding of sperm to zona pellucida, etc. Notably, oxidation-reduction process was also significantly enriched (Fig 3). 1,062 transcripts (including 22 lncRNAs) were found to be specifically down-regulated in day5_*ybx1*^*+/+*^ vs day5_*ybx1*^*-/-*^, which involve protein transport, glycolytic process, gluconeogenesis, etc. (Fig 3). Similarly, we have also detected many transcripts differentially expressed in day5_*ybx1*^*+/+*^ vs day6_*ybx1*^*+/+*^ but not present in day5_*ybx1*^*+/+*^ vs day5_*ybx1*^*-/-*^ (Table 1). Concretely, 342 transcripts (including 2 lncRNAs) exhibit up-regulation in day6_*ybx1*^*+/+*^ and functional analysis showed that the functions of those DE RNAs might involve oxidation-reduction process, transmembrane transport, proteolysis, etc. (Fig 3). 48 specific down-regulated RNAs were found in day6_*ybx1*^*+/+*^ compared to day5_*ybx1*^*+/+*^ and their corresponding functional analysis showed that there are no BP categories significantly enriched. These functional analyses based on the different subsets of DE transcripts indicated that reduced expression of Ybx1 could trigger up-regulation of oxidative stress-related biological processes and pathways [e.g., oxidation-reduction process was significantly enriched by specific up-regulated transcripts in day5_*ybx1*^*-/-*^ and day6_*ybx1*^*+/+*^]. These findings suggest that reduced expression of Ybx1, either caused by Ybx1 knockout or innate autoregulation, exhibits strong correlation with the redox regulation in zebrafish. Noteworthily, many DE lncRNAs are included in those DE RNA datasets (the majority of them is found in the comparison of day5_*ybx1*^*+/+*^ vs day5_*ybx1*^*-/-*^), implying specific lncRNAs present in redox regulation following Ybx1 knockout.

**Table 1.** Basic statistics of differentially expressed transcripts in two different comparisons.

| 5D WT vs 5D MT | 5D WT vs 6D WT | Number (mRNAs/lncRNAs) |
| --- | --- | --- |
| Up | Up | 138/0 |
| Down | Down | 5/0 |
| Down | Up | 7/0 |
| Up | Down | 2/0 |
| Up | Non | 709/22 |
| Down | Non | 1040/22 |
| Non | Up | 340/2 |
| Non | Down | 48/0 |

**Fig 3.**
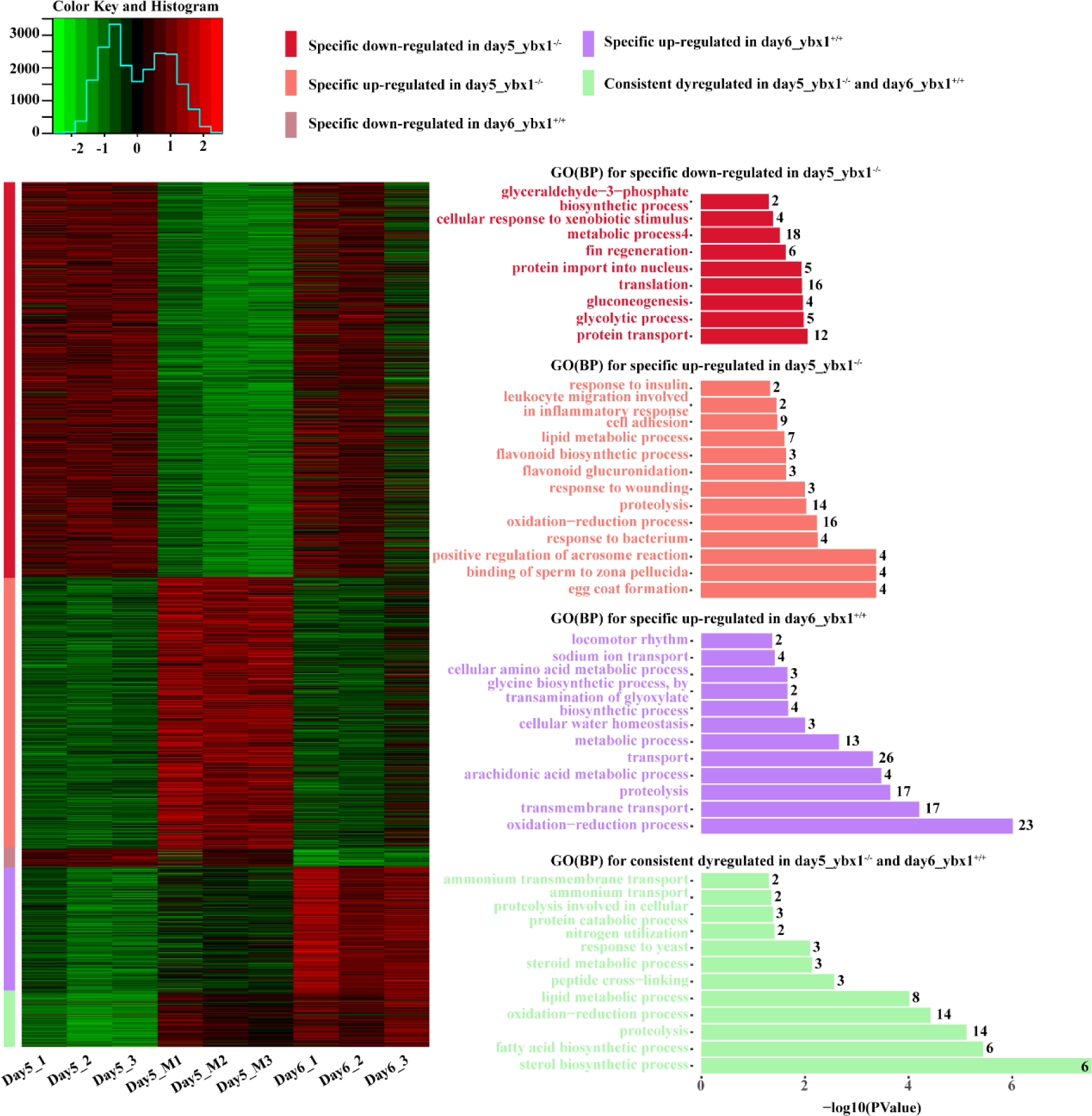
Differentially expressed transcripts and associated annotation terms. The heatmap shows transcript expression after normalization by DESeq2 for five different subsets of differentially expressed transcripts, including specific down-regulated in day5_ybx1^−/−^, specific up-regulated in day5_ybx1^−/−^, specific down-regulated in day6_ybx1^+/+^, specific up-regulated in day6_ybx1^+/+^ and consistent dyregulated in day5_ybx1^−/−^ and day6_ybx1^+/+^. Values have been centered and scaled for each row. Each row represents a single transcript. Statistically significant function annotation (BP) terms identified using DAVID (FDR<5%).

### Network analysis reveals lncRNAs implicated in Redox regulation

To investigate the lncRNAs as well as their possible roles implicated in Ybx1 knockout, a systematic co-expression network analysis was conducted using WGCNA in R (25). Initially, the flashClust tools package was used to conduct cluster analysis on these samples to detect outliers, the results showed that all samples are in the clusters and passed the cuts (Fig S5). Then a network-topology analysis was conducted for several soft-thresholding powers in order to have relative balance scale independence and mean connectivity of the WGCNA. The lowest power for the scale-free topology fit index was selected for construction of hierarchical clustering tree (Fig S6). This analysis co-localizes both correlated lncRNAs and mRNAs into 95 modules (Fig S7). Each module contains independent datasets of transcripts (Table 2). The interactions among those modules were visualized in Fig S8. The modules with common expression patterns that were significantly associated with specific traits (day5_*ybx1*^*-/-*^ and day6_*ybx1*^*+/+*^) were detected based on the correlation between module eigengene and traits. Then the function plotEigengeneNetworks of WGCNA was used to identify groups of correlated eigengenes. The results indicated that two modules (yellow and black) were significantly correlated to the traits: day5_*ybx1*^*-/-*^ and day6_*ybx1*^*+/+*^, respectively (Fig 4). The relationship of transcript significance with module membership was visualized in Fig S9. The yellow modules significantly related to day5_*ybx1*^*-/-*^ include 2,394 mRNAs and 352 lncRNAs, whereas black modules correlated to day6_*ybx1*^*+/+*^ contain 1,075 mRNAs and 155 lncRNAs (Table 2).

**Table 2.** The number of mRNAs and lncRNAs in the top 10 modules.

| Module | Size | lncRNA | mRNA |
| --- | --- | --- | --- |
| turquoise | 4731 | 318 | 4413 |
| blue | 4284 | 328 | 3956 |
| brown | 2752 | 382 | 2370 |
| yellow | 2746 | 352 | 2394 |
| green | 2267 | 355 | 1912 |
| red | 1563 | 176 | 1387 |
| black | 1230 | 155 | 1075 |
| pink | 1206 | 207 | 999 |
| magenta | 1109 | 98 | 1011 |
| purple | 970 | 201 | 769 |

**Fig 4.**
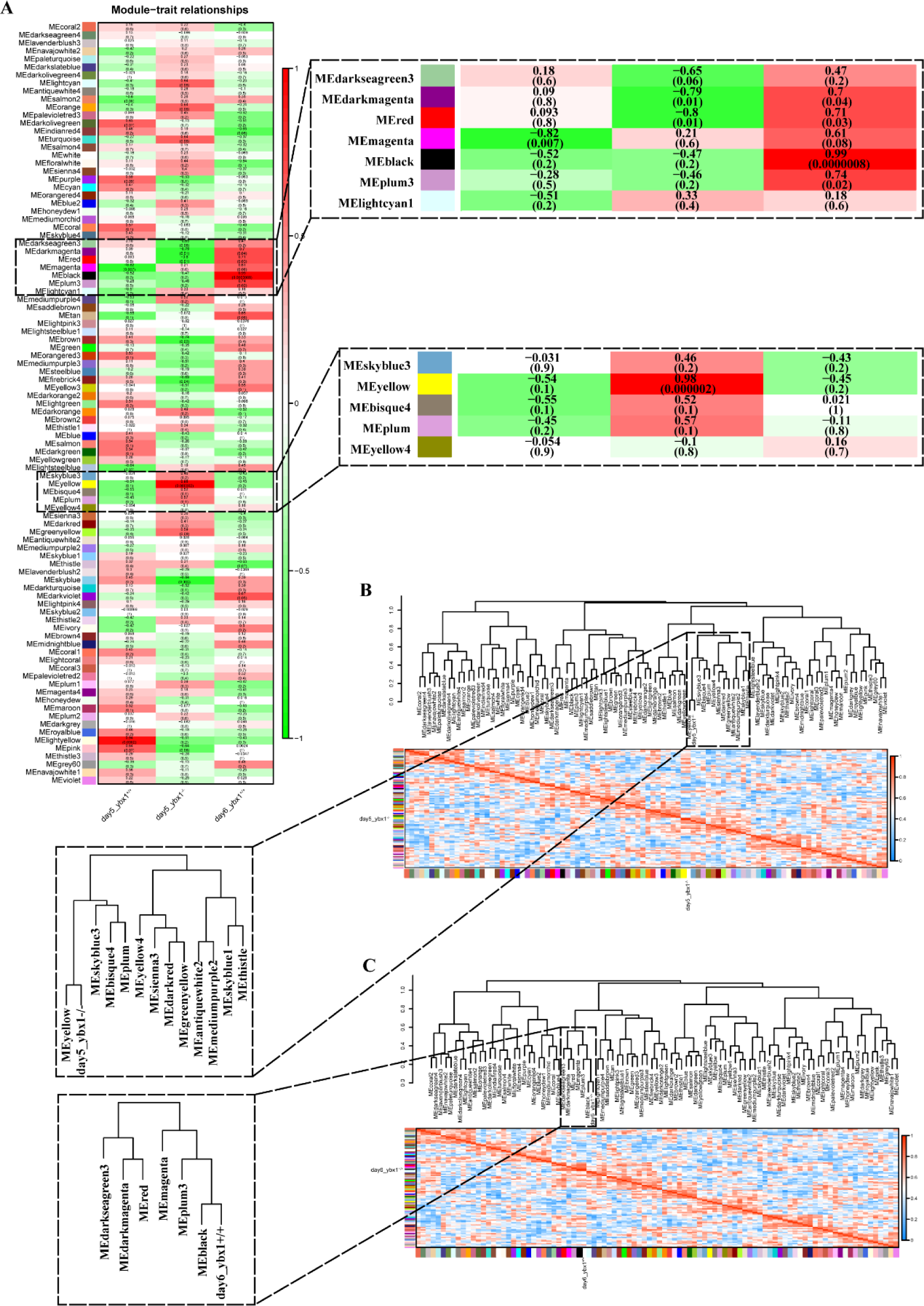
Detection of two trait-related modules. A) Module-trait associations. Each row corresponds to a module eigengene, column to a trait. Each cell displays the corresponding correlation and p-value. B) The eigengene dendrogram and heatmap 800 identify yellow module is highly related to day5_ybx1^−/−^. C) The eigengene dendrogram and heatmap identify black module is highly related to day6_ybx1^+/+^.

Go enrichment analyses based on the DAVID server were subsequently performed using the annotated transcripts (mRNAs) derived from two trait-related modules. Unsurprisingly, the results indicated that many Redox processes were significantly enriched by up-regulated transcripts in both two trait-related modules with considerably high significance (highlighted in red text in Fig 5), including GO:0006979 (response to oxidative stress), GO:0016491 (oxidoreductase activity) and GO:0055114 (oxidation-reduction process), etc. This finding is consistent with the results of functional enrichment analysis using DE RNAs described in the previous section (Fig 3), which further suggests the decline of Ybx1 can enhance up-regulation of Redox. Indeed, many studies (26–28) have illustrated the influence of oxidative stress to embryonic development in zebrafish, e.g., oxidative stress condition could change the cellular redox state, which further causes oxidation of molecules such as membrane peroxidation, loss of ions, protein cleavage, DNA strand breakages, and finally leads to cell death.

**Fig 5.**
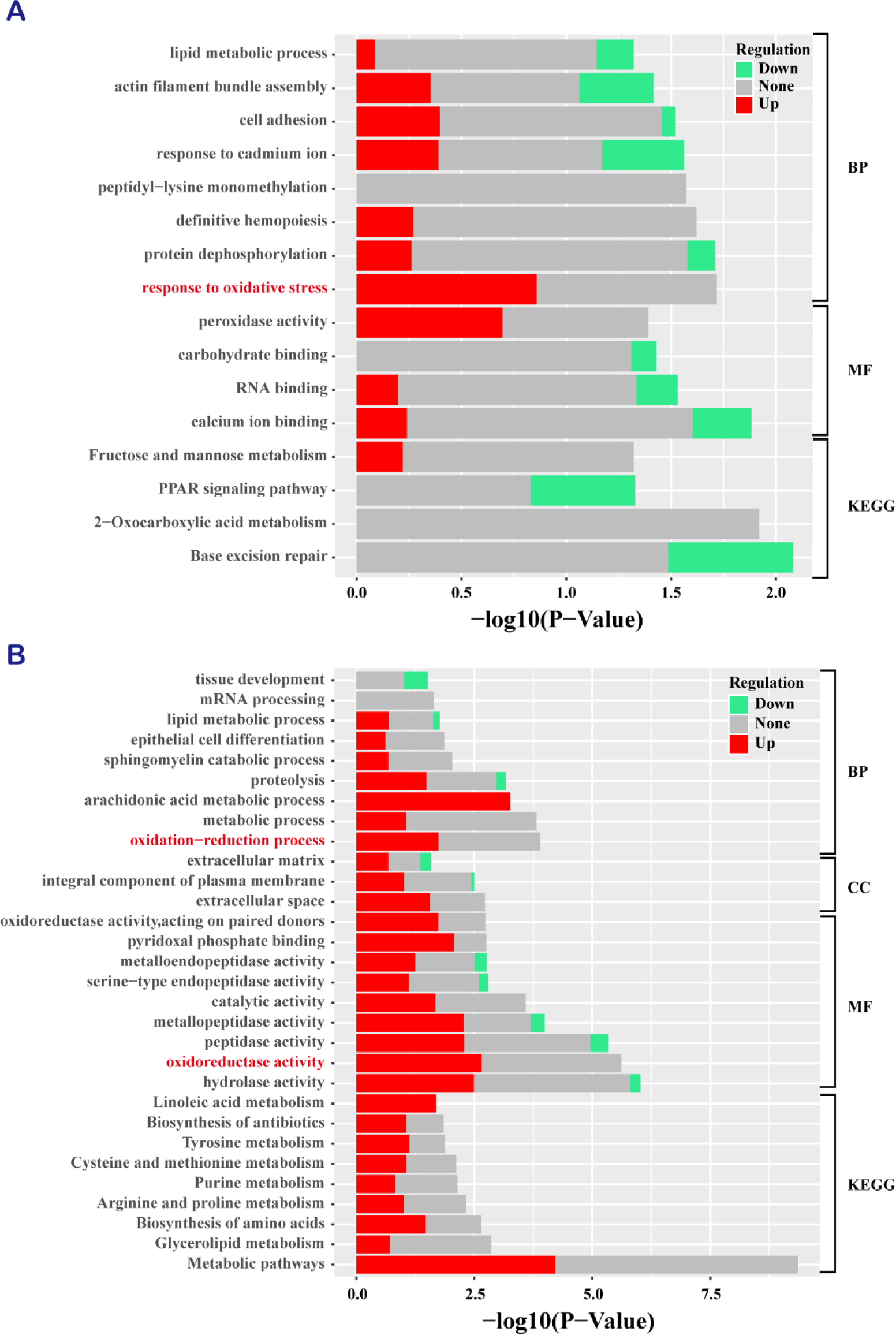
GO functional enrichment analysis of two trait-related modules: A) day5_ybx1^−/−^-related module. B) day6_ybx1^+/+^-related module. The colored bar reports the fraction of upregulated and downregulated transcripts present in that functional category. All the adjusted statistically significant values of the terms were negative 10-base log transformed.

Next, to explore possible roles of lncRNAs in Redox regulation triggered by decreased Ybx1, the expression profile of DE lncRNAs and DE mRNAs included in the yellow module related to day5_*ybx1*^*-/-*^ were used to assess the intramodular connectivity (including mRNA-mRNA and mRNA-lncRNA interactions) using TOM (Topology Overlap Matrix) similarity, which was done by the function *TOMsimilarityFromExpr* of WGCNA package. We did not further investigate the lncRNAs implicated in Redox in black module related to day6_*ybx1*^*+/+*^, since our previous DE analysis showed that a few DE lncRNAs (only 2) were found in the comparison of day5_*ybx1*^*+/+*^ vs day6_*ybx1*^*+/+*^. Regarding yellow module, we detected a great number of interactions which involved 631 DE mRNAs and 45 DE lncRNAs (Fig 6-A). Among them, several lncRNAs as well as mRNAs were found to interact with REDOX-related mRNAs, i.e., *duox* and *noxo1a* (Fig 6-B). These two genes have been demonstrated to play important roles in response to oxidative stress (29, 30). Also, our functional enrichment analysis indicated that both mRNAs were significantly enriched in the REDOX process, e.g., GO:0006979 (response to oxidative stress) and GO:0055114 (oxidation-reduction process). Hence, our subsequent work focused on the lncRNAs exhibiting correlation with *duox* and *noxo1a*. Particularly, we observed that several lncRNAs not only exhibiting strong correlations to Ybx1 but can also interact with two mRNAs that could be translated to two closely Redox-related proteins, Duox and Noxo1a (Fig 6-C). To test and confirm our findings that those specific lncRNAs associated to REDOX regulation in zebrafish embryonic development, we selected three lncRNAs, i.e., ENSDART00000171757, MSTRG30533.1 and MSTRG33365.1 exhibiting relatively significant differentially-expressed in day5_*ybx1*^*+/+*^ compared to day5_*ybx1*^*-/-*^, which simultaneously have strong correlations with Duox and Noxo1a as well as Ybx1 from the sub-network (Fig 6) for further experimental validation study.

**Fig 6.**
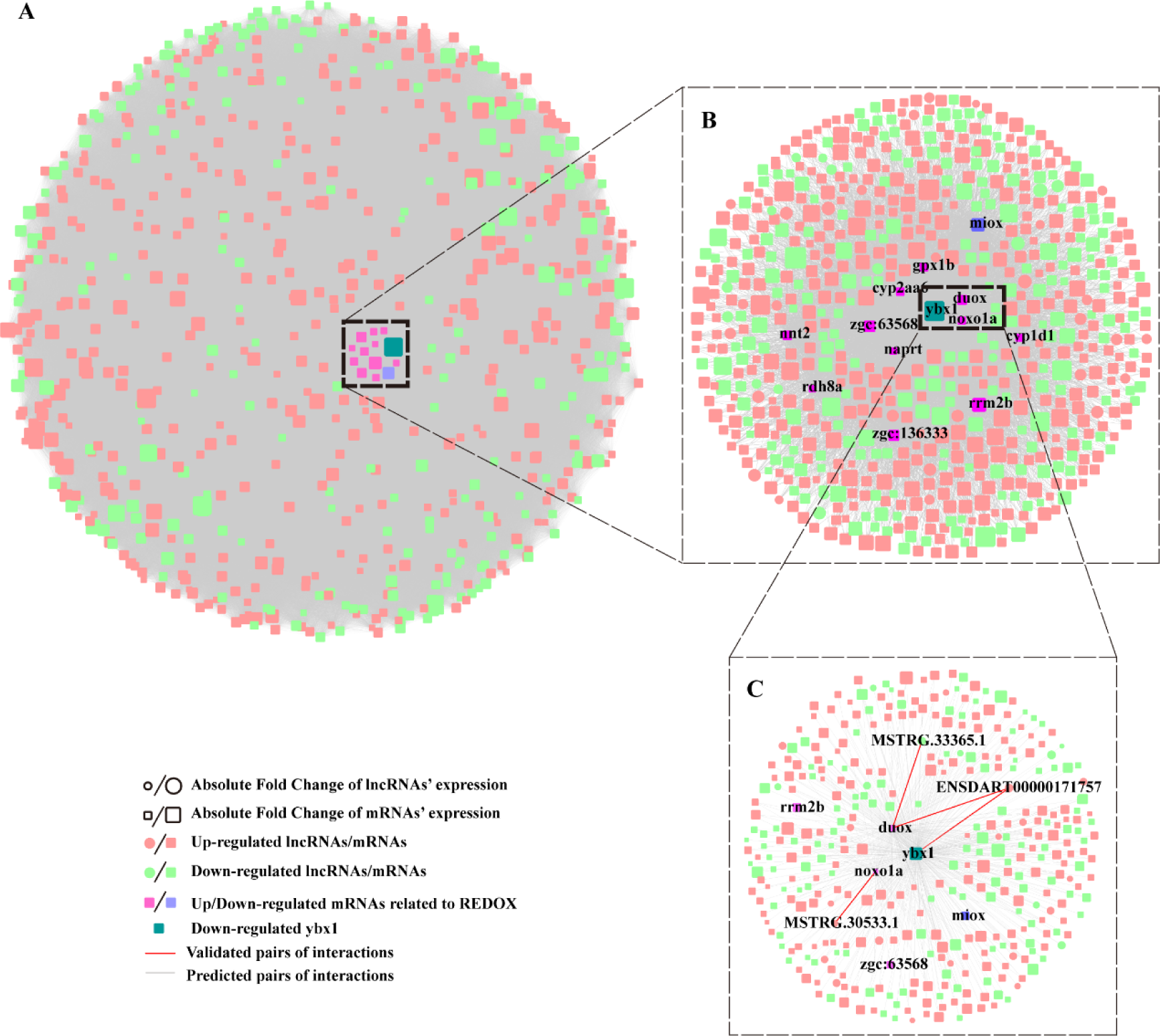
Co-expression network analysis defined REDOX-related interactome in day5_ybx1^−/−^-related module. A) Cytoscape map of the interaction (including mRNA-mRNA, mRNA-lncRNA and lncRNA-lncRNA) network defined by differentially expressed lncRNAs and mRNAs using the function *TOMsimilarityFromExpr* of WGCNA package. B) A zoomed view of a core network displaying interactions (including mRNA-mRNA and mRNA-lncRNA) involved in REDOX-related genes and YBX1. C) Zoomed in subnetwork view of interactions involved in two key REDOX-related genes: DUOX, NOXO1a, as well as YBX1.

### Differential expression of lncRNAs validation by RT-qPCR

First, we validated the differentially expressed lncRNAs using real-time qPCR. Apart from three selected lncRNAs, ENSDART00000171757, MSTRG30533.1, MSTRG33365.1 (which have been demonstrated as important lncRNAs related to Redox in the previous section), another two lncRNAs, MSTRG.12630.1, MSTRG24792.1 which have relatively high fold change in differential analysis between both day5_*ybx1*^*+/+*^ and day5_*ybx1*^*-/-*^ were selected for real-time qPCR validation. The expression levels of the five lncRNAs in day5_*ybx1*^*+/+*^ and day5_*ybx1*^*-/-*^larvae were exhibited in Fig 7. The expressions of MSTRG.12630.1, MSTRG24792.1, ENSDART00000171757, MSTRG30533.1 were all apparently increased in day5_*ybx1*^*-/-*^ larvae (Fig 7A-D), and MSTRG33365.1 presented sharp decrease in day5_*ybx1*^*-/-*^ larvae (Fig 7E). As expected, both the RNA-seq data and RT-qPCR results exhibited consistent expression patterns in day5_*ybx1*^*+/+*^ and day5_*ybx1*^*-/-*^ larvae.

**Fig. 7.**
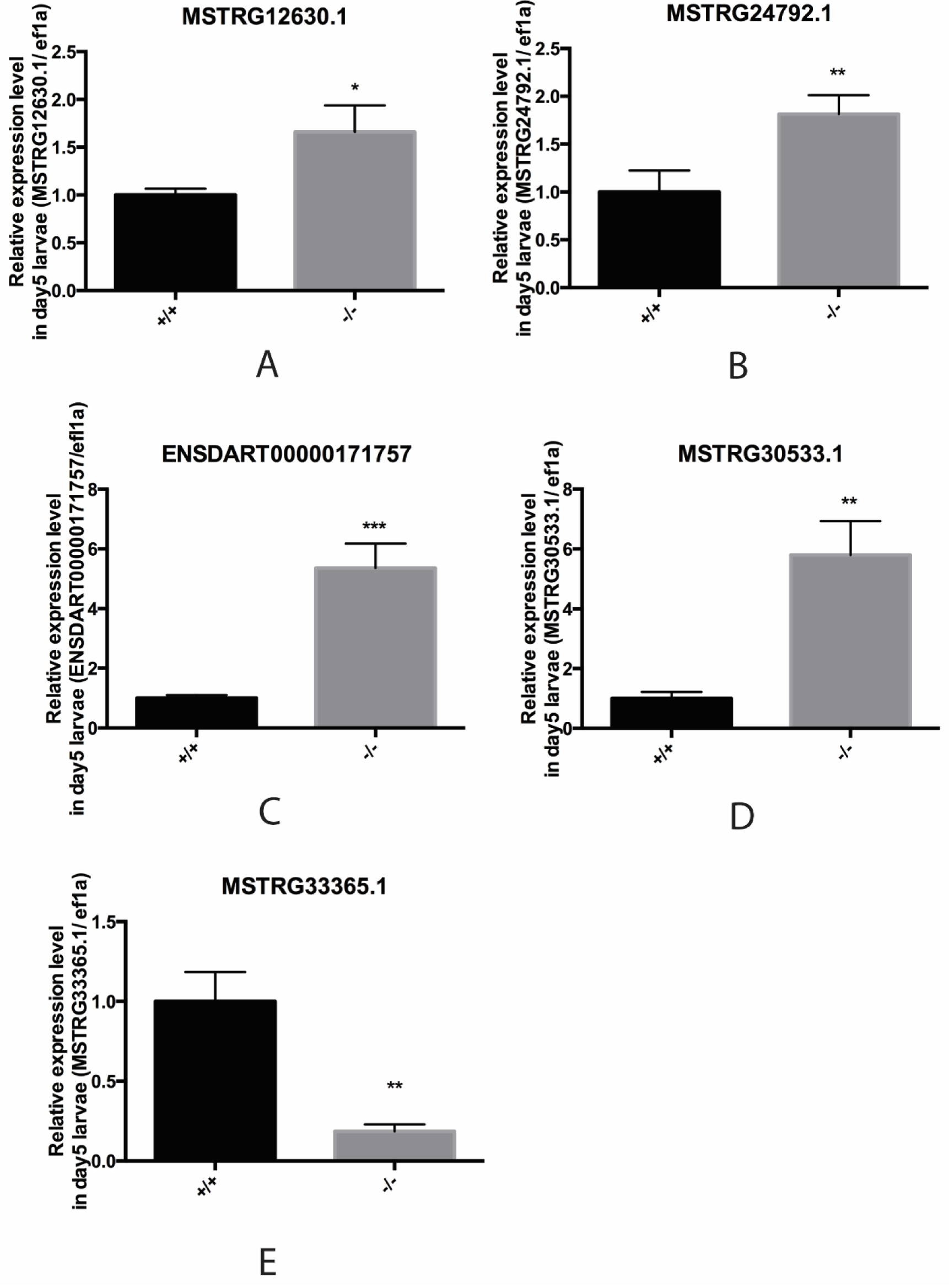
Reverse transcription and real-time quantitative PCR to validation of differentially expressed lncRNA identified by RNA-seq between day5 *ybx1* ^+/+^ and *ybx1* ^−/−^ larvae. A-E: Relative expression level of five selected differential lncRNA in both day5 *ybx1* ^+/+^ and *ybx1* ^−/−^ larvae, they are MSTRG12630.1, MSTRG24792.1, ENSDART00000171757, MSTRG30533.1, MSTRG33365.1 respectively. *:p value <0.05, **:p value<0.01, ***:p value<0.001.

### Spatial expression study reveals lncRNAs specifically expressed in gut

Firstly, we applied whole mount in situ hybridization (WISH) for spatial expression pattern study in both day5_*ybx1*^*+/+*^ and day5_*ybx1*^*-/-*^ larvae to test if the expression of those selected lncRNAs has tissue-specificity. By WISH assay, we discovered all the three lncRNAs were predominantly expressed in the gut region (Fig 8A-F). In addition, the difference of their expression levels between day5_*ybx1*^*+/+*^ and day5_*ybx1*^*-/-*^ larvae was quantified and was consistent with RT-qPCR results (Fig 8G-I).

**Fig. 8.**
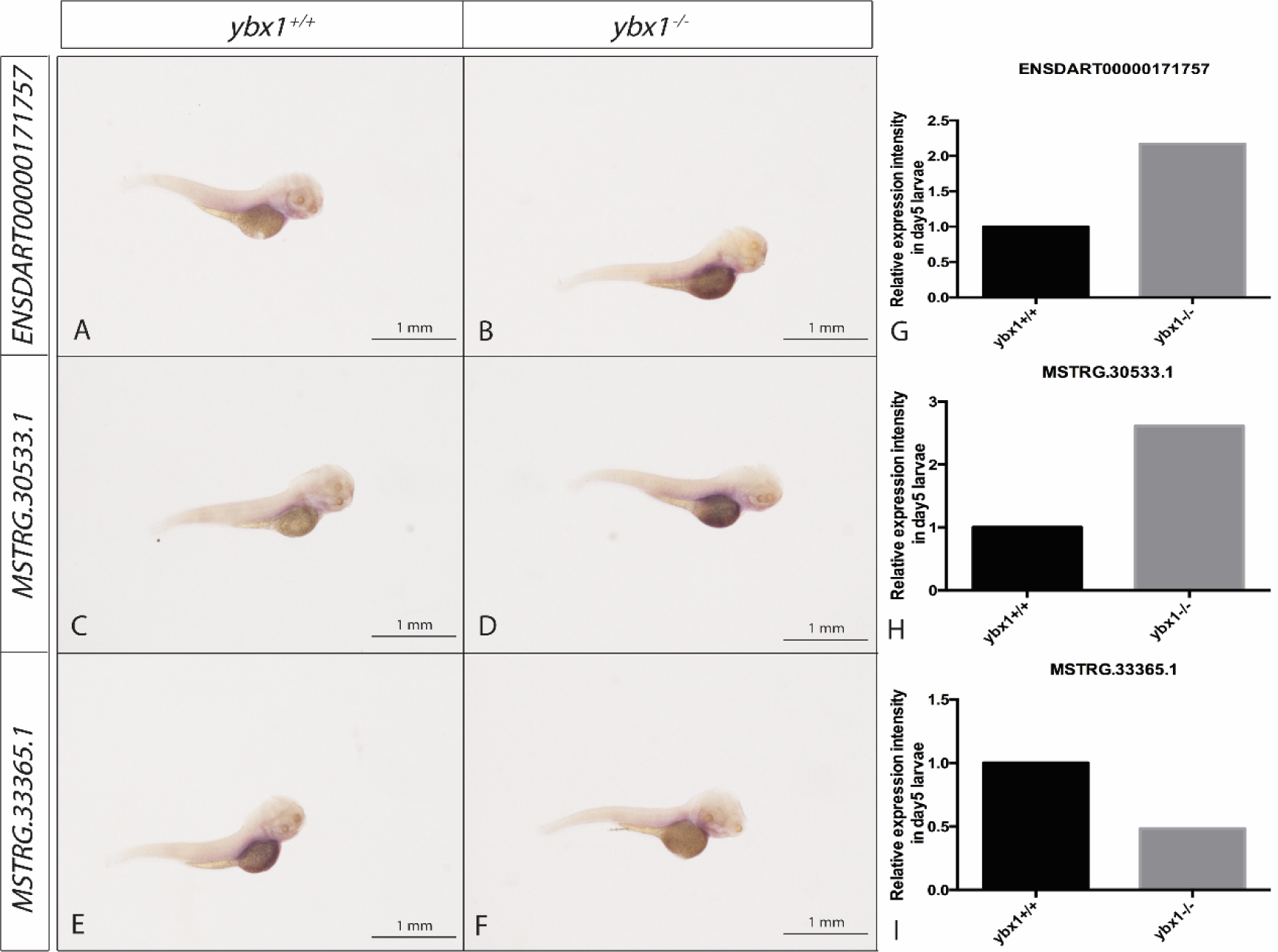
Whole mount in situ hybridization (WISH) analysis of selected three lncRNA (ENSDART00000171757, MSTRG.30533.1, MSTRG.33365.1) in day5 *ybx1* ^+/+^ and *ybx1* ^−/−^ larvae. A-B: the expression of ENSDART00000171757 lncRNA in day5 *ybx1* ^+/+^ and *ybx1* ^−/−^ larvae. C-D: the expression of MSTRG.30533.1 lncRNA in day5 *ybx1* ^+/+^ and *ybx1* ^−/−^ larvae. E -F: lncRNA MSTRG.33365.1 expression in day5 *ybx1* ^+/+^ and *ybx1* ^−/−^ larvae. G-I: Relative expression intensity of ENSDART00000171757, MSTRG30533.1, MSTRG33365.1 in day5 *ybx1* ^+/+^ and *ybx1* ^−/−^ larvae.

### lncRNAs knockdown lead to larvae morphological deformation

Our network analysis indicated that all three lncRNAs exhibiting strong correlation to Ybx1, which has been proven to be a key protein to zebrafish embryonic development. Therefore, it is reasonable to infer that these lncRNAs probably play a role in embryonic development. To test it, the three selected lncRNAs, ENSDART00000171757, MSTRG30533.1, MSTRG33365.1, were targeted for morpholino knockdown, with their specific target sequence for morpholino synthesis (Fig 9A). After serial morpholino injection (5ng, 10ng, 20ng), partial larvae on day5 with severe morphological deformation were photographed (Fig 9C). Interestingly, we discovered a large proportion of deformed larvae for each of the three lncRNAs morpholino, but only a few deformed larvae by control morpholino injection. In particular, the rate of larval deformation after morpholino injection was dose-dependent for all the three lncRNAs (Fig 9B). In conclusion, by comparison of the deformation rate between control morpholino and the three targeted morpholino, our experiments showed that the knockdown of each of the three selected lncRNAs leads to larvae morphological deformation in dose-dependent manner.

**Fig. 9.**
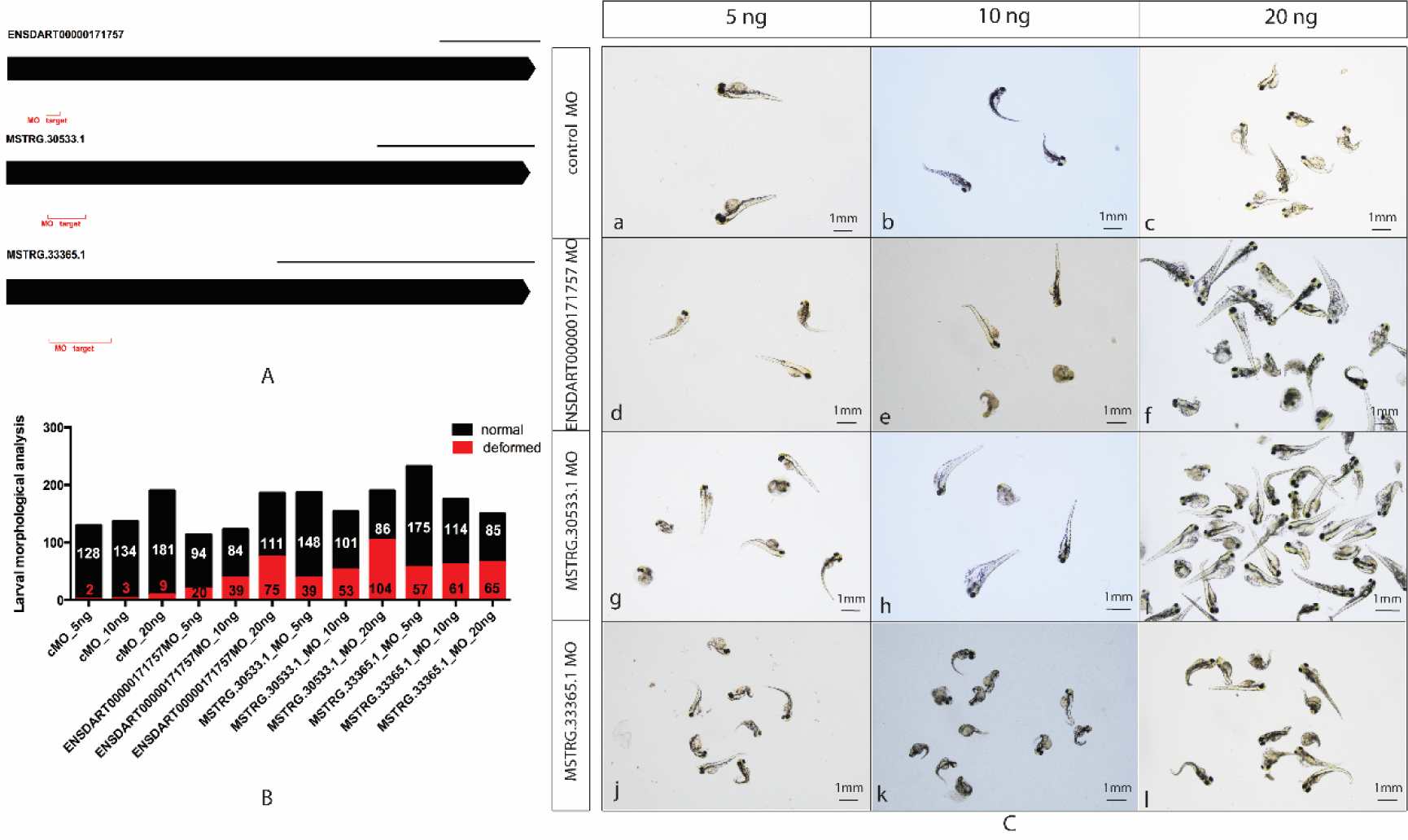
Larval morphological analysis after lncRNA morpholino injection on day5. A: Targeted sequence designed for specific morpholino synthesis for three lncRNA, ENSDART00000171757, MSTRG.30533.1, MSTRG.33365.1. B: Larval morphological statistics for different lncRNA morpholino injected larvae with different doses (5ng, 10ng, 20ng) injection, control morpholino injection was used for reference. Column in black represents larvae with normal morphology, and red column stands for deformed morphological larvae. C: Photographs of deformed larvae injected with different doses (5ng, 10ng, 20ng) of morpholino for lncRNA (ENSDART00000171757, 846 MSTRG.30533.1, MSTRG.33365.1), and control morpholino injected larvae were used 847 as reference.

### Correlation study reveals lncRNAs-mediated interactions in Redox regulation

Our network analysis also indicated that three selected lncRNAs exhibit strong correlations to the Redox-related genes *duox* and *noxo1a*, i.e., 327 ENSDART00000171757-*duox*, MSTRG.30533.1-*noxo1a*, MSTRG.33365.1-*duox*. To study their interactions, larvae (day5) injected with 20ng morpholino for the three lncRNAs were divided into two groups, normal morphology and deformed morphology. Knockdown of ENSDART00000171757 slightly reduced the mRNA expression of *duox* in both the normal and deformed groups (Fig 10A), and remarkably reduced the mRNA expression of *ybx1* in normal group (Fig 10B). MSTRG30533.1 knockdown apparently activated *noxo1a* mRNA expression in normal group (Fig 10C), while inhibition of MSTRG33365.1 led to a sharp decrease of *duox* mRNA expression in deformed group (Fig 10D). Briefly, our analysis demonstrated that both lncRNAs ENSDART00000171757 and MSTRG.33365.1 might positively regulate the expression of *duox*, while MSTRG30533.1 probably exhibits negative regulation to *noxo1a*. LncRNAs ENSDART00000171757 also exhibited positive regulation to Ybx1.

**Fig. 10.**
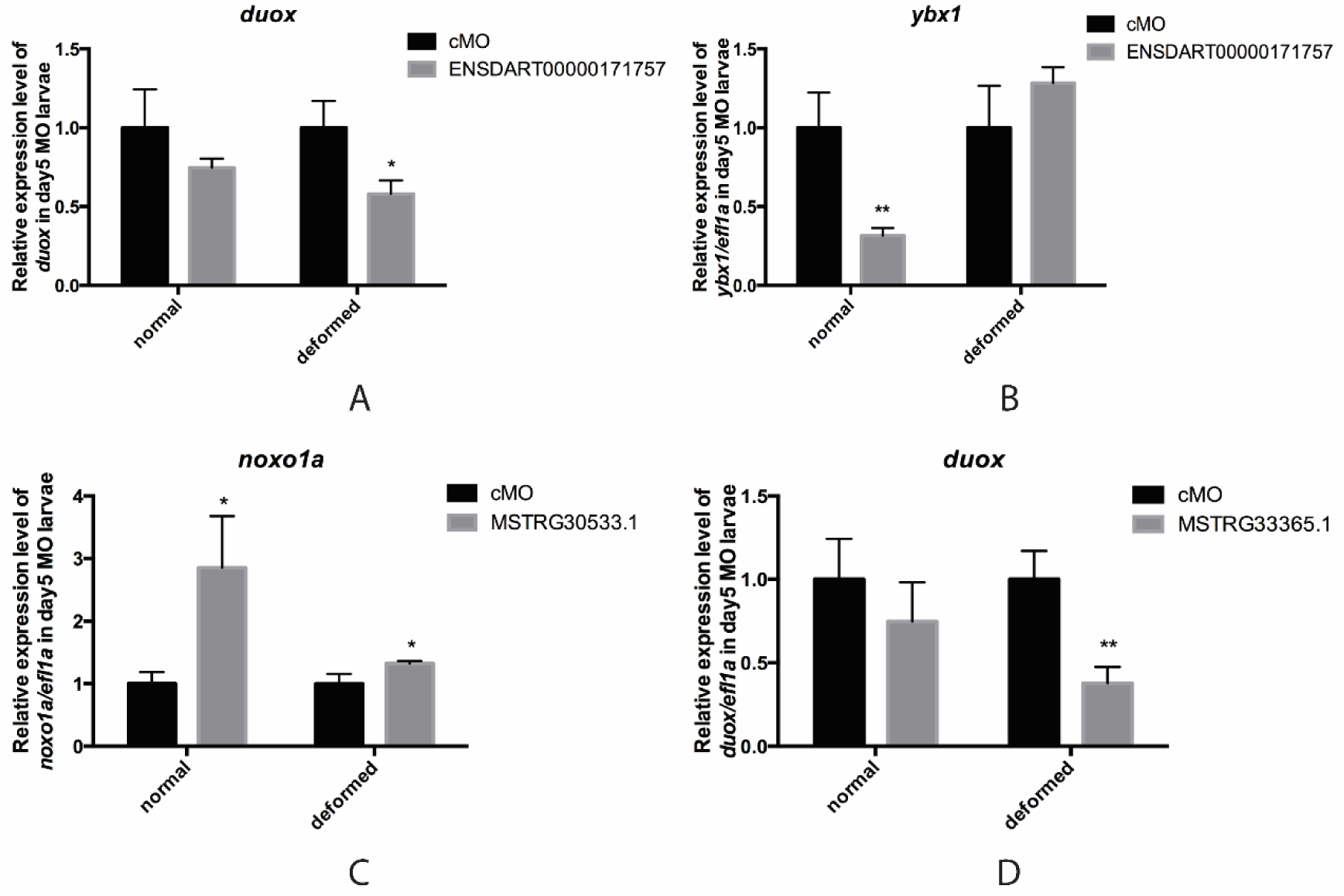
Reverse transcription and real-time quantitative PCR to study the correlation between selected lncRNA and potential targeted mRNA after targeted lncRNA morpholino injection. A-B: Relative expression of *duox* mRNA and *ybx1* mRNA in day5 larvae after injection of 20ng ENSDART00000171757 morpholino. C: Relative expression of *noxo1a* mRNA in day5 larvae after injection of 20ng MSTRG.30533.1 855 morpholino. D: Relative expression of *duox* mRNA in day5 larvae after 20ng MSTRG.33365.1 morpholino injection. 20ng control morpholino injected larvae were used as reference. On day5, morpholino injected larvae were divided into two groups, normal and deformed morphology. Column in black represents cMO, and grey column stands for targeted lncRNA morpholino. *: p value< 0.05, **: p value< 0.01.

Also, we further tested the potential interaction between ENSDART00000171757 and Ybx1 in protein level. Immunoblotting was performed to evaluate Ybx1 protein expression level under ENSDART00000171757 morpholino knockdown and beta-actin was used as internal reference. The results showed that the expression of Ybx1 protein was evidently reduced after ENSDART00000171757 knockdown (Fig 11A). It was suggested that inhibition of ENSDART00000171757 would result in significant reduction (approximately 50%) of the relative intensity of Ybx1/Actin in day5 larvae (Fig 11B). In conclusion, this experimental study well demonstrated the correlation between ENSDART00000171757 and Ybx1.

**Fig. 11.**
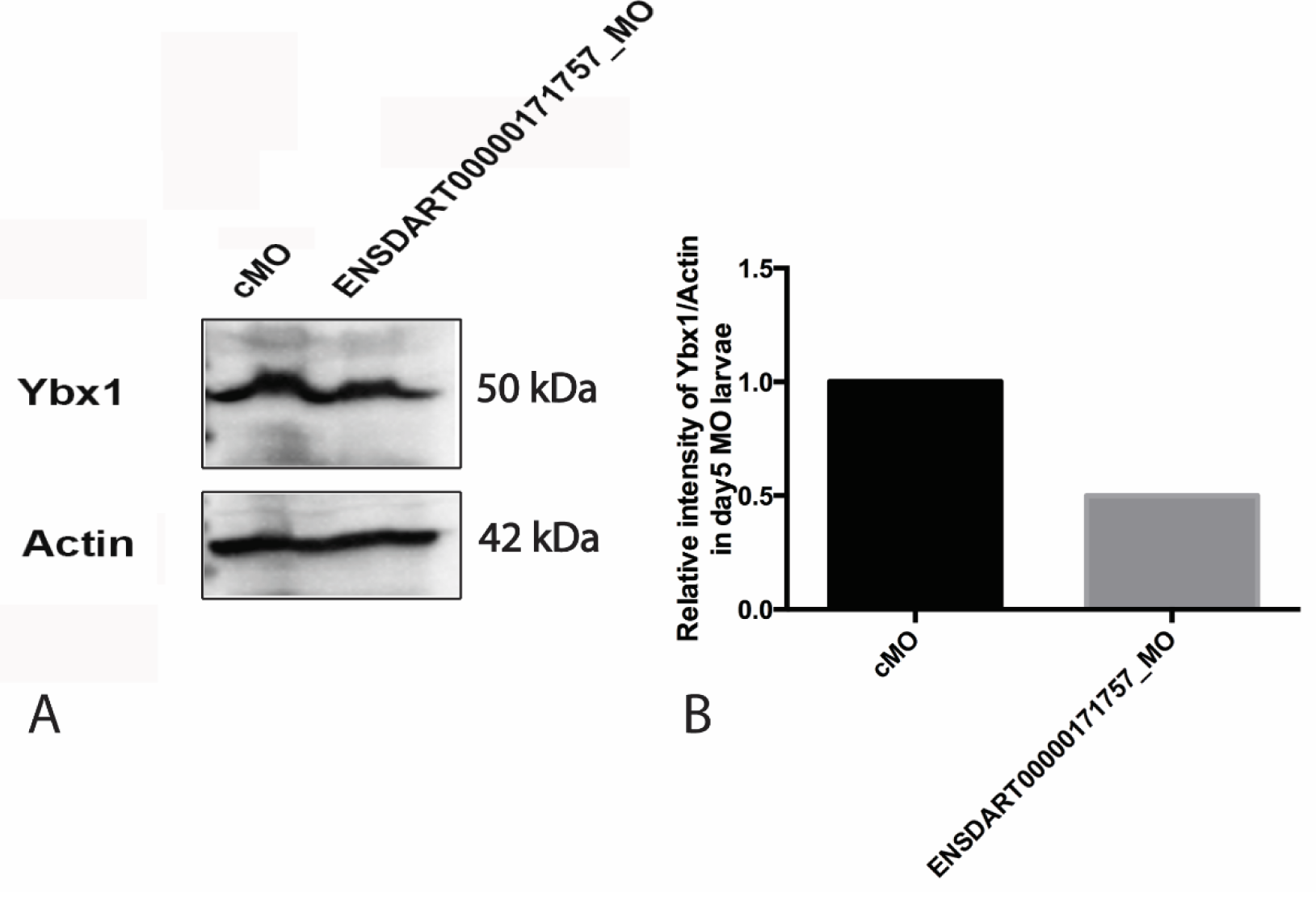
Immunoblotting study for validation of the correlation between Ybx1 expression and ENSDART00000171757 lncRNA knockdown. A: western blotting results of Ybx1 between cMO and ENSDART00000171757_MO (20ng) injected day5 larvae samples, larvae homogenized were with normal morphology. Beta-Actin expression level was used as reference. B: Relative intensity of Ybx1/Actin in day5 ENSDART00000171757_MO injected larvae, cMO injected larvae were as the control.

## Discussion

Given that lncRNAs have been proposed to carry out diverse functions, for instance, regulation of gene expression by acting as signaling, guiding, sequestering or scaffolding molecules, an increasing number of studies focuses on the disclosure of the roles of lncRNA in vertebrate development (31–33). In some cases, however, similar works on the same lncRNAs come to contradictory conclusions. A typical example is that Lin et al’s study (34) and Ulitsky et al’s study (35) showed that knockdown of the lncRNA megamind led to zebrafish embryonic defects, whereas Kok et al ‘s study (36) demonstrated that megamind had no influence on embryonic development. These inconsistent results have caused a dispute about whether lncRNA is important to vertebrate development. Notably, Mehdi Goudarzi1’s study (14) used CRISPR/Cas9 system to knockout 24 specific lncRNAs in zebrafish which was picked out based on synteny, conservation, expression dynamics and proximity to development-related genes and concluded that individual lncRNAs have no overt functions in zebrafish embryogenesis, viability and fertility. Given these facts, it is likely that lncRNAs exert limited functions in vertebrate development, particularly in zebrafish embryonic development. Under such research background, our study regards Ybx1 as the breakthrough of study point, utilizes TALENs technique to establish *ybx1* homozygous mutants (day5_*ybx1*^*-/-*^) in zebrafish and chooses two wild-type zebrafish strains (day5_*ybx1*^*+/+*^, day6_*ybx1*^*+/+*^) as a control group. Then with the aim of unveiling the existence of the lncRNAs implicated in zebrafish embryonic development, as well as their possible roles in it and relevance of Ybx1, a systematic transcriptome analysis was conducted based on deep RNA sequencing using Illumina platform. Ybx1 has been demonstrated as a curial gene for all stage of embryonic development in mouse and zebrafish (7, 8). Knockout of Ybx1 in zebrafish undoubtedly provides a good animal mmodel for our study.

Worthy of a special mention is the fact that despite the RNA library in the present study is prepared using oligo-dT to capture sequences with poly-A tail, a great amount of lncRNAs could still be sequenced. The reason is that lncRNAs are structurally similar to mRNAs in some way, i.e., the majority of mature lncRNAs is produced by the same RNA polymerase II transcriptional machinery and can be polyadenylated, suggesting some of them are indistinguishable from mRNAs. As a result, we initially reconstructed the zebrafish transcriptome, and then identified high-confidence dataset of lncRNAs using a stringent filtering pipeline. Next, to figure out the lncRNAs implicated in zebrafish embryonic development, we performed differential expression analysis and systematic network analysis between day5_*ybx1*^*-/-*^, day6_*ybx1*^*+/+*^ and day5_*ybx1*^*+/+*^. The aforesaid series of analysis concluded that decreasing of Ybx1 no matter by acquired knockout or innate autoregulation exhibits closely correlations to Redox processes which involved several lncRNA-mRNA interactions.

Embryo development is a complicated process that underlies precise regulation and control. The roles of Redox to embryonic development have been well elucidated so far. Concretely, oxygen can be regarded as a double-edged sword since it is essential to embryogenesis but also acts a potential hazard via formation of reactive oxygen and reactive nitrogen species (ROS/RNS) (37). Overmuch ROS leads to oxidative stress and further causes either cell death or senescence by oxidation of cellular molecules (38). Redox balance is thus critical to embryonic development. More importantly, ROS generated by nicotinamide adenine dinucleotide phosphate (NADPH) oxidases is a crucial redox signal to establish homeostasis (29). This knowledge directs us narrow down the huge number of mRNA-lncRNA interactions to a relatively small sub-network exhibiting strong correlation to two REDOX-related genes, *duox* (NADPH oxidase) and *noxo1a* (NADPH oxidase organizer). Due to the importance of these two genes to Redox regulation and relevance of embryo development, it can be inferred that the lncRNAs correlated to these two Redox-related genes also are important to embryo development in zebrafish. To test our prediction, we picked up three lncRNAs for validation bioassay. The experimental results indicated that all three lncRNA knockdown by morpholino led to deformation on dose-dependent manner in zebrafish larvae, implying their importance to zebrafish embryonic development. Furthermore, correlation study between lncRNAs and two Redox genes demonstrated their close associations. Above all, Ybx1 is an important protein to embryogenesis and development, disclosure of the correlation between Ybx1 and lncRNAs transcriptome will shed new insights for unveiling organelle-based mechanisms of lncRNAs implicated in zebrafish embryo development.

## Materials and methods

### Animals

The wild type zebrafish (*Danio rerio*) AB strain was used in this study. Zebrafish were maintained in ZebTEC multilinking rack system (Tecniplast; Buguggiate, Italy) under an artificial photoperiod of 14 h light:10 h dark. The temperature, pH and conductivity of the system were 28 ±1 °C, 7.5 and 400 µS/cm, respectively. The fish were fed twice a day with Otohime fish diet (Marubeni Nisshin Feed, Tokyo, Japan) by Tritone automatic feeding system (Tecniplast).

All experiments were performed under a license from the Government of Macau Special Administrative Region (SAR) and approved by Animal Experimentation Ethics Committee of the University of Macau.

### Larvae RNA preparation and RNA sequencing

Larvae from wild type zebrafish of AB strain (day5_*ybx1*^*+/+*^, day6_*ybx1*^*+/+*^) and *ybx1* homozygous mutants (day5_*ybx1*^*-/-*^) which were established by transcription activator-like effector nucleases (TALENs) were collected for larval RNA preparation. Around 15-20 zebrafish larvae of each type were collected and pooled for sufficient amount of RNA per sample. A total of 9 samples, day5_*ybx1*^*+/+*^ (3 duplicates), day6_*ybx1*^*+/+*^ (3 duplicates), day5_*ybx1*^*-/-*^ (3 duplicates), were included in RNA sequencing.

Total RNA was extracted using Tri-Reagent (Molecular Research Center, Cincinnati, OH) according to the protocol of the manufacturer and our previous report (39). Total RNA was then treated with DNase for 10 min at 37 °C to remove genomic DNA [10 μg RNA in 100 μl reaction buffer with 2U DNase I from NEB (Ipswich, MA)] followed by phenol-chloroform extraction and ethanol precipitation.

### Basic RNA-seq data analysis

Initially, the raw data were processed using Trimmomatic v0.36 (ILLUMINACLIP: TruSeq3-PE.fa:2:30:10:8:true SLIDINGWINDOW:4:15 LEADING:3 TRAILING:3 MINLEN:50) to remove low quality reads and adaptors. Clean reads were aligned against zebrafish genome (Ensembl release 91) (20) using STAR v020201 (16) with 2-pass mode. Then the alignment results were assembled into transcripts using StringTie v1.3.3b (17) with gene annotation reference. Finally, the assembled transcripts from each sample were merged into a consensus of final transcripts. The quality of the assembled transcripts was evaluated using REF-EVAL from DETONATE (v1.11) (18).

### lncRNAs identification

To identify lncRNAs, we applied a stringent stepwise filtering pipeline, which has been widely used in our previous studies (40, 41). At first, the assembled transcripts were annotated and excluded by aligning against known protein sequences from NCBI nr database, Uniprot database (42), and the mRNA and protein datasets of zebrafish derived from Ensembl database using BLAST (v2.6.0+) (19). This step aims at removing protein-coding sequences as much as possible. Then the remaining unannotated transcripts with length larger than 200 nt and longest ORF less than 100 residues were retained. In addition, Coding Potential Calculator (CPC) (21) was used in the second trimming of protein-coding sequences. Finally, the remaining transcripts were translated (stop-to-stop codon) using an in-house perl script and then subjected to Pfam database (43) to search for sequences probably encoding protein domains or motifs. The unmatched transcripts were retained as final high-confidence dataset of lncRNAs.

### Differential expression analysis

The assembled transcripts in each sample were quantified by featurecounts (v1.5.3) (44). Then the expression profile was normalized using median of ratios method by R package DESeq2 (45). Differential expression analysis was also performed using DESeq2. Differentially expressed (DE) RNAs were obtained by cutoff of adjusted P-value<0.05 (Benjamin-Hochberg correction) (45).

### Functional prediction of lncRNAs based on network analysis

In order to investigate the possible functions of lncRNAs implicated in Ybx1 knockout, a weighted co-expression network analysis was performed based on the differential expressed lncRNAs and mRNAs using a package WGCNA in R (25). Prior to WGCNA analysis, the expression profiles of lncRNAs and mRNAs were normalized using regularized log normalization in DESeq2. To construct the network and detect the modules related to traits (day5_*ybx1*^*-/-*^ and day6_*ybx1*^*+/+*^), the function *softConnectivity* from WGCNA was used with the “randomly selected genes” parameter set at 5000 and the power parameter pre-calculated by pickSoft-Threshold function of WGCNA. Then the predicted modules of highly correlated RNAs were subjected to Metascape to search for relevant biological processes and pathways. The intramodular connectivity of mRNA and lncRNA in trait-related modules was assessed using TOM similarity, which were calculated by *TOMsimilarityFromExpr* of WGCNA. TOM similarity reflects the potential interactions between transcripts.

### Reverse Transcription and Real-Time Quantitative PCR (RT-qPCR)

Reverse transcription was performed at 37° C for 2 h in a total volume of 10 μl reaction solution containing 3 μg RNA, 0.5 μg oligo (dT), 1X MMLV RT buffer, 0.5 mM each dNTP, 0.1 mM dithiothreitol, and 100 U M-MLV reverse transcriptase (Invitrogen, Carlsbad, CA). To validate the RNA-seq data, we determined the expression levels of several selected lncRNA transcripts by RT-qPCR in both day5_*ybx1*^*+/+*^ and day5_*ybx1*^*-/-*^, including MSTRG12630.1, MSTRG24792.1, ENSDART00000171757, MSTRG.30533.1, MSTRG.33365.1. To study the correlation between the selected lncRNA and their targets, including *duox, noxo1a*, and *ybx1*, the mRNA expression levels of targets were determined by RT-qPCR after targeted lncRNA morpholino microinjection. The expression levels were normalized to that of the housekeeping gene *ef1a*. The standard for each gene was prepared by PCR amplification of cDNA fragments with specific primers (Table 3). The real-time qPCR assay was performed on the CFX96 Real-time PCR Detection System (Bio-Rad, Hercules, CA) and repeated twice. All values were expressed as the mean ± SEM (*n* = 3), and the data were analyzed by *t*-test using Prism 6 on Macintosh OS X (GraphPad Software, San Diego, CA).

**Table 3.**
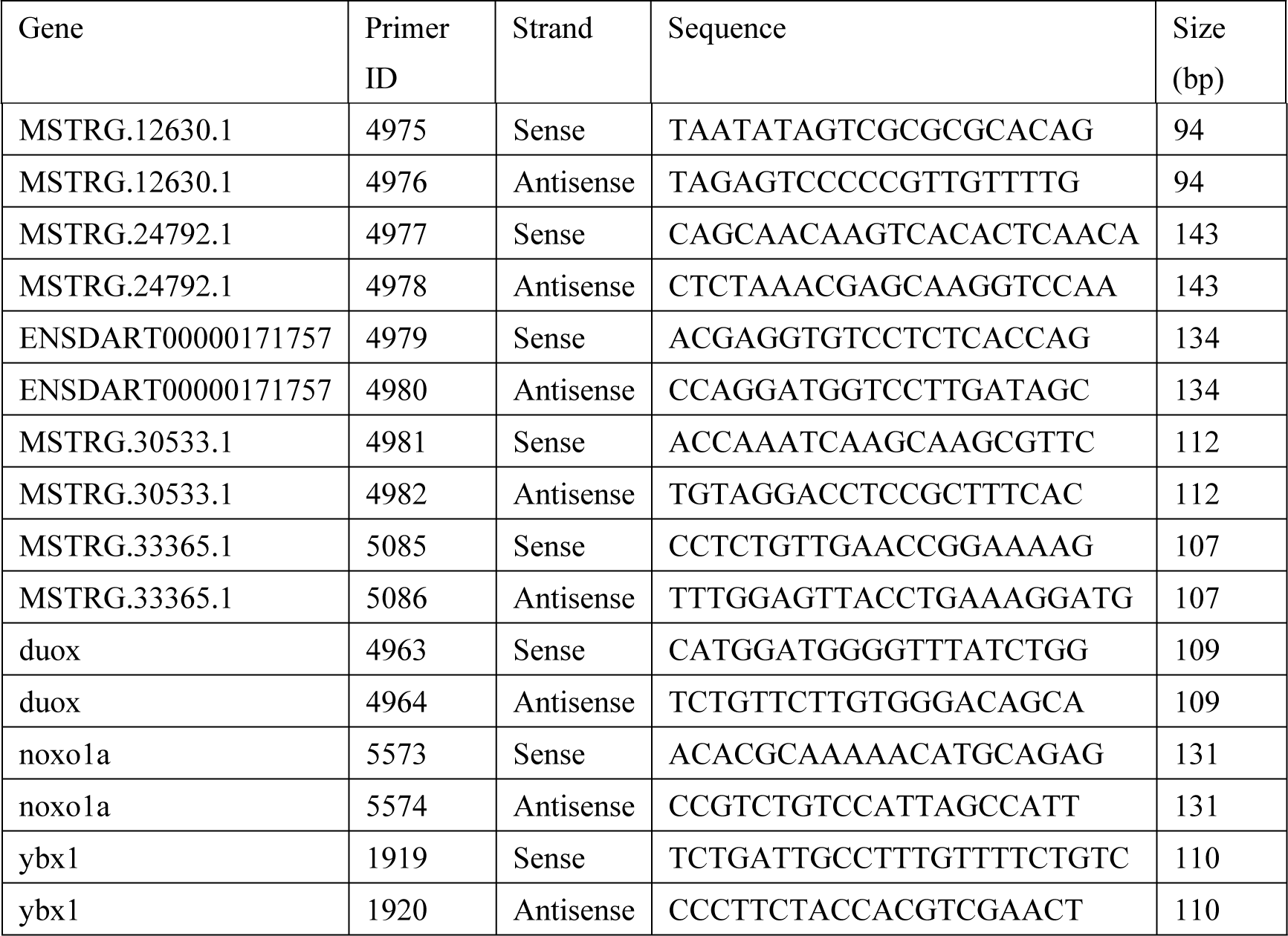
Sequence information of primers for RT-qPCR.

### Morpholino knockdown and morphological analysis

To investigate the functions of three selected lncRNA, ENSDART00000171757, MSTRG.30533.1, MSTRG.33365.1, 300 nmol morpholino and 100nmol standard morpholino oligo were synthesized by Gene Tools (Gene Tools, Philomath, OR). The morpholino targeted sequences for synthesis regarding to lncRNA above were represented in Table 4. For morpholino microinjection, three different doses (5ng, 10ng, 20ng) of morpholino were injected into one cell stage embryos for targeted lncRNA respectively within 30 minutes just after spawning. Equal amount (5ng, 10ng, 20ng) of standard morpholino oligo was injected as control. On day5, count the embryo number both with normal and deformed morphology individually for different morpholino and different dose injection. The morphology of morpholino injected larvae were photographed by Nikon SMZ18 microscope (Nikon, Tokyo, Japan).

**Table 4.**
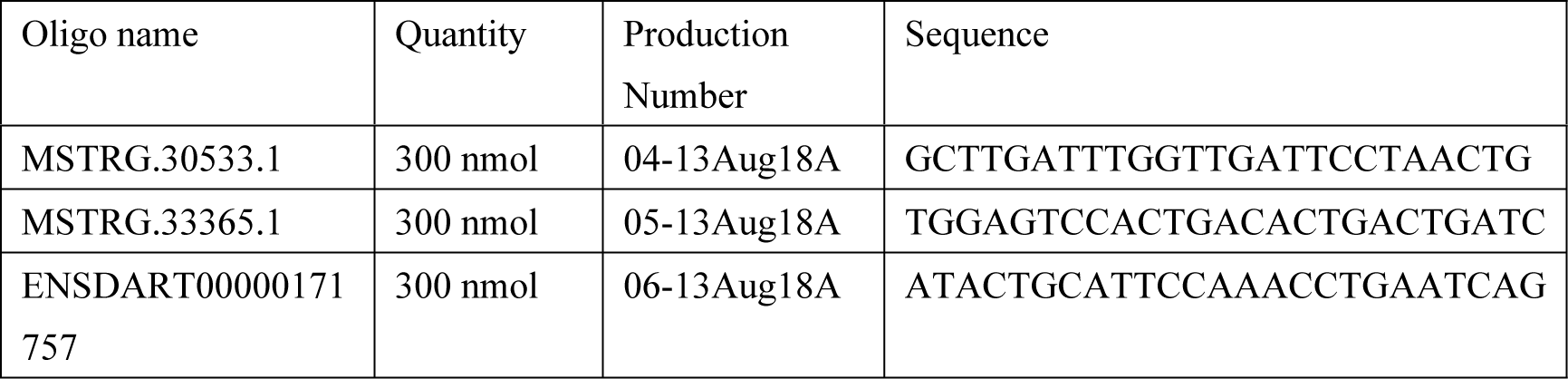
Morpholino information for three lncRNA.

### Immunoblotting

To further study the correlation between lncRNA ENSDART00000171757 and *ybx1* expression at protein level, western blotting was used for evaluation of Ybx1 protein expression between morpholino (ENSDART00000171757) injected and control morpholino injected groups. The larvae on day 5 with normal morphology were collected for protein sample preparation.

Briefly, larvae were lysed with 100 µL 1x SDS sample buffer (62.5 mM Tris-HCl, 1% w/v SDS, 10% glycerol, 5% mercaptoethanol, pH=6.8), and the lysates were heated at 95 °C for 10 minutes and then centrifuged 12,000 rpm for 15 minutes at 4 °C and followed by protein concentration measurement. The samples (total 100 µg proteins) from ENSDART00000171757_MO and control MO were separated on 12% polyacrylamide gels and transferred to PVDF membranes. The membranes were blocked with 5% non-fat milk in 1x TBST for 1 h at room temperature. After washing three times with 1x TBST, the membranes were incubated with Ybx1 primary antibody (1:5000) and beta-actin primary antibody (1:2000) in 5 ml of 1x TBST with 5% non-fat milk overnight at 4 °C. They were washed with 1x TBST three times followed by incubation with the HRP-conjugated secondary antibody (1:2000) in 5 ml 1x TBST for 1 h at room temperature. After washing, the membranes were incubated with the SuperSignal West Femto Maximum Sensitivity Substrate (Thermo Scientific, Waltham, MA) and signals detected on the ChemiDoc MP imaging system (Bio-Rad, Hercules, CA).

### Whole mount in situ hybridization

To further study the spatial expression pattern for the three target lncRNA, ENSDART00000171757, MSTRG.30533.1, MSTRG.33365.1, whole mount RNA in situ hybridization (WISH) were undertaken for investigation. All their lncRNA probes were transcribed in vitro and for storage at −80°C. Embryos were under 0.003% 1-phenyl-2-thiourea (PTU) treatment to remove pigment until the larvae reach to day5. On day 5, the *ybx1*^*+/+*^ and *ybx1*^*-/-*^ larvae were fixed in fresh 4% PFA overnight at 4°C. Wash larvae with PBST for 5 minutes at room temperature, and repeat 4 times. For permeabilization, the larvae were immersed with 10 µg/ml Protease K in PBST for 15 minutes at room temperature. The larvae were refixed in 4% PFA for 20 minutes at room temperature, followed by PBST rinsing for 5 minutes that was repeated 4 times. Before mRNA probe hybridization, larvae were immersed in hybridization mix for 2 hours at 65°C. Dilute 100 ng labelled RNA probe in hybridization mix. Remove prehybridization solution and add pre-warmed hyb mix plus probe to the larvae. The hybridization was kept for 40 hours at 65°C. For post-hybridization washes, wash 5 min in 66% hybridization mix, 33% 2x SSC at 65°C, and wash 5 min in 33% hybridization mix, 66% 2x SSC at 65°C, wash 5 min in 2x SSC at 65°C. Wash 1x 20 min in 0.2x SSC+0.1% Tween-20 at 65°C, and then wash 2x 20 min in 0.1x SSC+0.1% Tween-20 at 65°C. At room temperature, wash 5 min in 66% 0.2x SSC, 33% PBST, and wash 5 min in 33% 0.2x SSC, 66% PBST, wash 5 min in PBST. After washing, incubate larvae in blocking solution (PBST plus 2% sheep serum, 2 mg/ml BSA) 1 hour at room temperature, prepare first antibody (alkaline-phosphatase conjugated anti-digoxigenin) by diluting it in blocking solution; 1:5000. Incubate in antibody for shaking overnight at 4°C, and wash 5×15 min in PBST. For colorization, wash 4×5 min in coloration buffer. Mix 200 µL nitro-blue tetrazolium (NBT)/5-bromo-4-chloro-3-indolyl phosphate (BCIP) stock mixture with 10 ml coloration buffer. Add 500 µL of this mix to larvae and incubate in the dark at room temperature until a blue reaction product is visible. Stop reaction by quickly washing larvae 3 times by stop solution (PBST, pH 5.5), then washes twice in stop solution for 15 minutes. After glycerol mounting, photograph immediately by Nikon SMZ18 microscope (Nikon, Tokyo, Japan). 569

## Supporting information

**Table S1 Basic statistics of zebrafish RNA-seq data before and after quality trimming**.

(DOCX)

**Table S2 Basic statistics of assembly results of transcriptome in zebrafish**.

(DOCX) 577

**Table S3. Basic statistics of quality assessment of assembled transcripts achieved by DETONATE**.

(DOCX)

**Fig S1. Heatmap of differentially expressed transcripts detected in comparison between day5_ybx1**^**+/+**^ **vs day5_ybx1**^**-/-**^.

(DOCX)

**Fig S2. Volcano Plot of differentially expressed transcripts detected in comparison between day5_ybx1**^**+/+**^ **vs day5_ybx1**^**-/-**^. **Each spot represents a transcript. Red spots represent differentially expressed transcripts**.

(DOCX)

**Fig S3. Heatmap of differentially expressed transcripts detected in comparison between day5_ybx1**^**+/+**^ **vs day6_ybx1**^**+/+**^.

(DOCX)

**Fig S4. Volcano Plot of differentially expressed transcripts detected in comparison between day5_ybx1**^**+/+**^ **vs day6_ybx1**^**+/+**^. **Each spot represents a transcript. Red spots represent differentially expressed transcripts**.

(DOCX)

**Fig S5. Sample clustering to detect outliers. All the samples were in the clusters, all samples have passed the cuts**.

(DOCX)

**Fig S6. Analysis of network topology for various soft-thresholding powers. The left panel indicates the scale-free fit index (y-axis) as a function of the soft-thresholding power (x-axis). The right panel shows the mean connectivity (degree, y-axis) as a function of the soft-thresholding power (x-axis)**.

(DOCX)

**Fig S7. Clustering dendrograms of transcripts, with dissimilarity based on topological overlap, together with assigned module colors**.

(DOCX)

**Fig S8. Visualizing the gene network using a heatmap plot. Light color represents low overlap and progressively darker red color represents higher overlap**.

(DOCX)

**Fig S9. Scatterplots of Gene Significance (GS) for recurrence vs Module Membership (MM) in the yellow module (A) and black module (B)**.

(DOCX)

## Acknowledgements

This work was funded by University of Macau through Research Grants MYRG2018-00071-FHS, EF005/FHS-ZXH/2018/GSTIC and FHS-CRDA-029-002-2017.

## Competing interests

All authors declare that no competing interests exists.

